# A cell-matrix interaction regulates the undifferentiated state and self-renewal capacity of avian primordial germ cells

**DOI:** 10.1101/2025.01.28.635251

**Authors:** Kennosuke Ichikawa, Yuzuha Motoe, Tenkai Watanabe, Ryo Ezaki, Mei Matsuzaki, Hiroyuki Horiuchi, Michael J. McGrew

**Affiliations:** The Roslin Institute, Royal (Dick) School of Veterinary Studies, University of Edinburgh, Midlothian, EH25 9RG, United Kingdom; Graduate School of Integrated Sciences for Life, Hiroshima University, 1-4-4 Kagamiyama, Higashi-Hiroshima, Hiroshima 739-8528, Japan

**Keywords:** Primordial germ cells, Chicken, Epithelial-to-mesenchymal transition, Somatic lineage conversion

## Abstract

Primordial germ cells (PGCs) are lineage restricted precursor cells of sperm and eggs. Whilst avian PGCs from chicken can be cultured and modified to produce genome-edited chickens, the long-term culture of PGCs from other bird species has not been achieved. Here, we explored the effects of a cell-matrix interaction on the in vitro propagation of chicken PGCs. Blocking integrin signaling severely reduced the self-renewal of the PGC, indicating that a PGC-matrix interaction is an essential process for self-renewal. We investigated the properties of somatic cell differentiated PGCs and expression analyses suggested that the PGCs undergo a partial epithelial-to-mesenchymal transition caused by excess PGC-matrix interactions. Finally, we conducted a long-term culture of chicken PGCs with matrix components, resulting in significant induction of their somatic conversion. These results suggested that a high level of PGC-matrix interaction can cause somatic conversion, whilst a moderate level is essential for their proliferation. Overall, we identified a molecular aspect of self-renewal and maintaining undifferentiated states of avian PGCs.

## 1. Introduction

Primordial germ cells (PGCs) are lineage restricted embryonic stem cells which give rise to the future gametes. They form early during avian development and migrate to colonize the gonadal anlagen. Currently, chickens (*Gallus gallus*) are the only vertebrate species whose PGCs can be stably cultured for an extended period [1, 2]. Genome-edited chickens have been widely produced through gene modification and transplantation of cultured PGCs [3]. Additionally, cultured PGCs can be cryopreserved and used for the conservation of local and commercial chicken breeds [4].

Expanding germ cell culture to non-chicken avian species will facilitate genome editing and the study of disease transmission in other bird species. For example, ducks (*Anas platyrhynchos*), the natural host of the avian influenza virus [5], are an essential experiment organism for studies on avian host immune responses [6–8]. Also, quails (*Coturnix japonica*) are an excellent experimental model owing to their short generation time and their value in studies of the developmental biology [9]. In addition, a PGC culture system may aid in the conservation of endangered avian species [10]. However, to date, stable long-term culture of PGCs from non-chicken avian species, such as ducks [11] and quails [12], remains unachieved.

Understanding the molecular mechanisms enhancing a self-renewal capacity with maintaining the undifferentiated state of cultured PGCs is essential for these processes. Here, we focused on a somatic fate decision process of avian PGCs through cell-matrix interactions. Avian PGCs, which are propagated in vitro as semi-adherent suspension cells, occasionally spontaneously differentiate into adherent somatic cells [13]. This phenomenon implies a matrix interaction triggers their conversion into a somatic cell linage. Contradictorily, a recent study showed avian PGCs themselves express matrix components, such as collagens and fibronectin, and a significant level of PGC-matrix interaction occurs during early embryogenesis [14]. In vitro propagation of chicken PGCs in low calcium medium can inhibit the conversion into a somatic cell linage and enhance stable growth [2], but somatic conversion and poor self-renewal capacities of PGCs are hypothesized to be a barrier to the long-term culture of PGCs retaining germ cell characteristic in non-chicken avian species.

In this study, we first reveal that inhibition of integrin signals reduces the self-renewal capacity of chicken PGCs, indicating that a PGC-matrix interaction is an essential factor for in vitro propagation. Next, we characterized chicken PGCs undergoing somatic cell differentiation, which was correlated with an induction of PGC-matrix interactions. Expression analyses and gene expression profiling by RNA-sequencing (RNA-seq) analyses suggests that PGCs convert to a somatic cell linage through a partial epithelial-to-mesenchymal transition (EMT) in accordance with activation of integrin signaling and alteration of expression patterns of integrins. We further reveal that the long-term culture of PGCs with matrix components significantly induced a somatic conversion, suggesting that a robust cell-matrix interaction triggers this conversion to somatic cell lineages. These results indicated that moderate levels of PGC-matrix interaction are required for their self-renewal, but excess integrin signals disrupt their undifferentiated states.

## 2. Results

### 2.1. The effect of a cell-matrix interaction on PGC self-renewal

We evaluated whether a cell-matrix interaction is required for the proliferation of the PGCs, as they are generally cultured as semi-adherent suspension cells on tissue culture plates [1, 15, 16]. Here, we used a low-calcium (0.15 mM) medium [2] optimized for the stable culture of chicken PGCs. We first cultured PGCs in an agarose gel coated well, which inhibits cell adhesion to the well surface [17]. We observed that the number of PGCs cultured in agarose coating well was significantly reduced (Fig. 1A-C). We next investigated Focal adhesion kinase (FAK) signaling, the major downstream kinase pathway activated by integrin interactions [18]. Culturing the PGCs with defactinib, an inhibitor of FAK phosphorylation, dramatically reduced the self-renewal capacity of the PGCs (Fig. 1D-F). Western blot analysis showed that PGCs cultured with DMSO (negative control) expressed phosphorylated-(phospho-) FAK, whilst the defactinib reduced the level of the phospho-FAK (Fig. 1G & Supplementary Fig. S1). These results suggested that activation of the FAK signaling through a PGC-matrix interaction is essential for PGC self-renewal in culture.

**Fig. 1.**
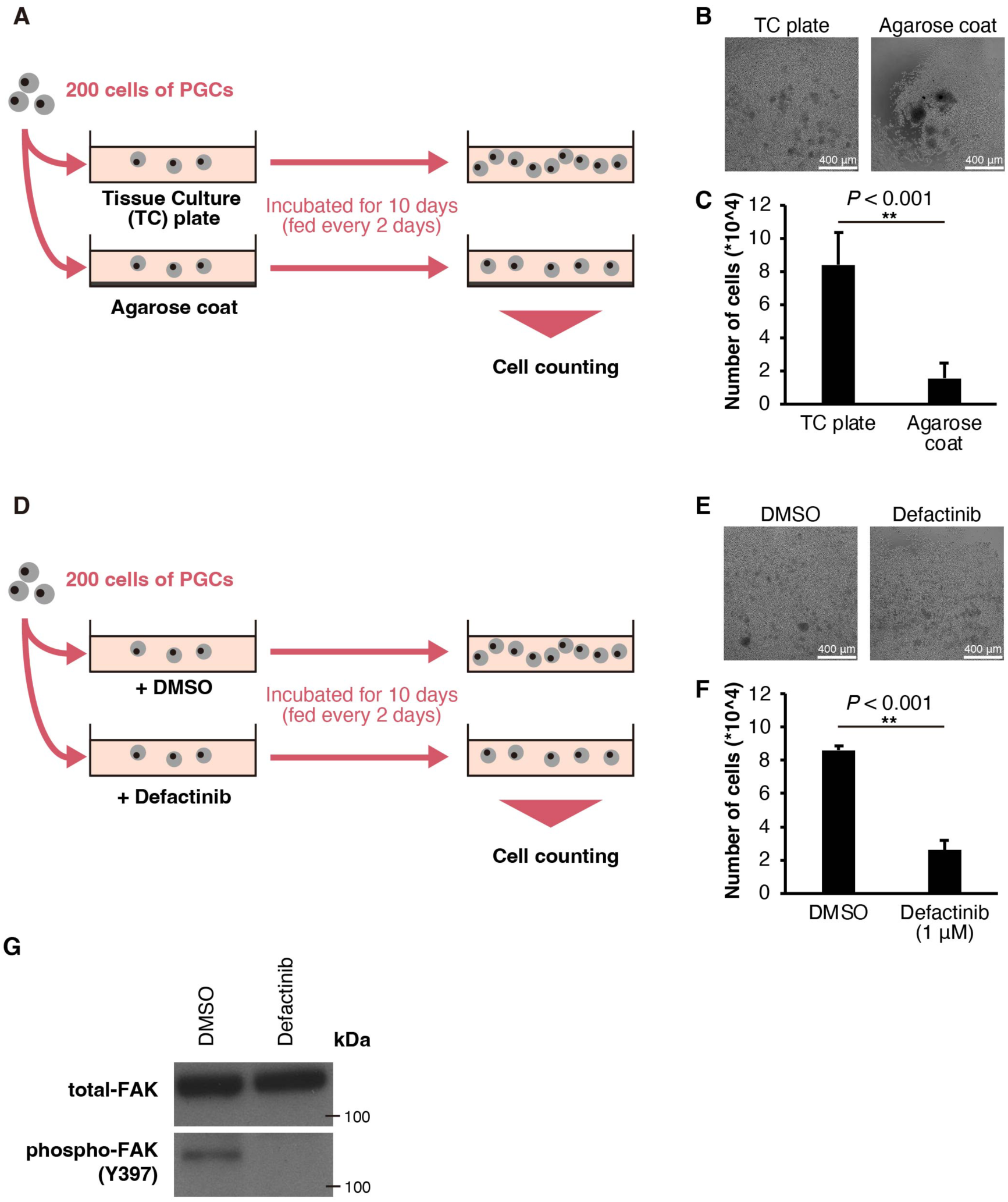
Evaluation of effects of integrin signaling on PGC self-renewal. (A) A schematic diagram of the experiment using TC plates and agarose coated plates. (B) Images of PGCs cultured for 10 days in TC plates or agarose coated plates. (C) The number of cells seeded at 200 cells/well and subsequently incubated for 10 days in each condition. Error bars indicate SD of the mean (n = 4 technical replicates). Statistical significance in each condition was evaluated by the *t*-test. Asterisks represent significant differences (\*\**P* < 0.01). (D) A schematic diagram of the experiment using DMSO and defactinib. (E) Images of PGCs cultured for 10 days with 0.1 % DMSO or 1 μM of defactinib. (F) The number of cells seeded at 200 cells/well and subsequently incubated for 10 days in each condition. Error bars indicate SD of the mean (n = 3). Statistical significance in each condition was evaluated by the *t*-test. Asterisks represent significant differences (\*\**P* < 0.01). (G) Western blot of treated and control cells. Molecular weight markers are indicated to the right of the membranes. Representative blot (n = 2). Uncropped images of each blot are shown in the Supplementary Fig. S1.

### 2.2. Characterization of somatic differentiated chicken PGCs focusing on germ cell markers

We next characterized in vitro propagated chicken PGCs undergoing spontaneous differentiation into somatic cells (Fig. 2A), which we hypothesized to be induced by a PGC-matrix interaction. We used a physiological normal (1.8 mM CaCl_2_) medium [19] in order to increase the PGC-matrix interactions and somatic cell conversion. We also used a Matrigel coating to enable them to adhere to the surface of chamber slides (Fig. 2B). We first measured the expression of the PGC marker, alkaline phosphatase (AP), during this process. AP-positive cells reflect undifferentiated states of both PGCs and ES cells, and previous studies showed that chicken PGCs are AP-positive [20, 21]. 48 hours after seeding the PGCs on Matrigel, we observed that adherent PGCs displayed two cellular morphologies. PGCs displaying a fibroblast-like morphology were AP-negative, whereas PGCs retaining a round morphology were AP-positive (Fig. 2C). To verify the somatic cell conversion of the fibroblast-like adherent PGCs, we performed immunofluorescence staining for DEAD-box helicase 4 (DDX4), a pan-germ cell marker [22]. Adherent PGCs with a fibroblast-like morphology were DDX4-negative, whereas PGCs retaining a round morphology were DDX4-positive (Fig. 2D). These initial results indicate cell morphology changes and germ cell/ pluripotency marker down regulation accompanies differentiation to the somatic cell lineage.

**Fig. 2.**
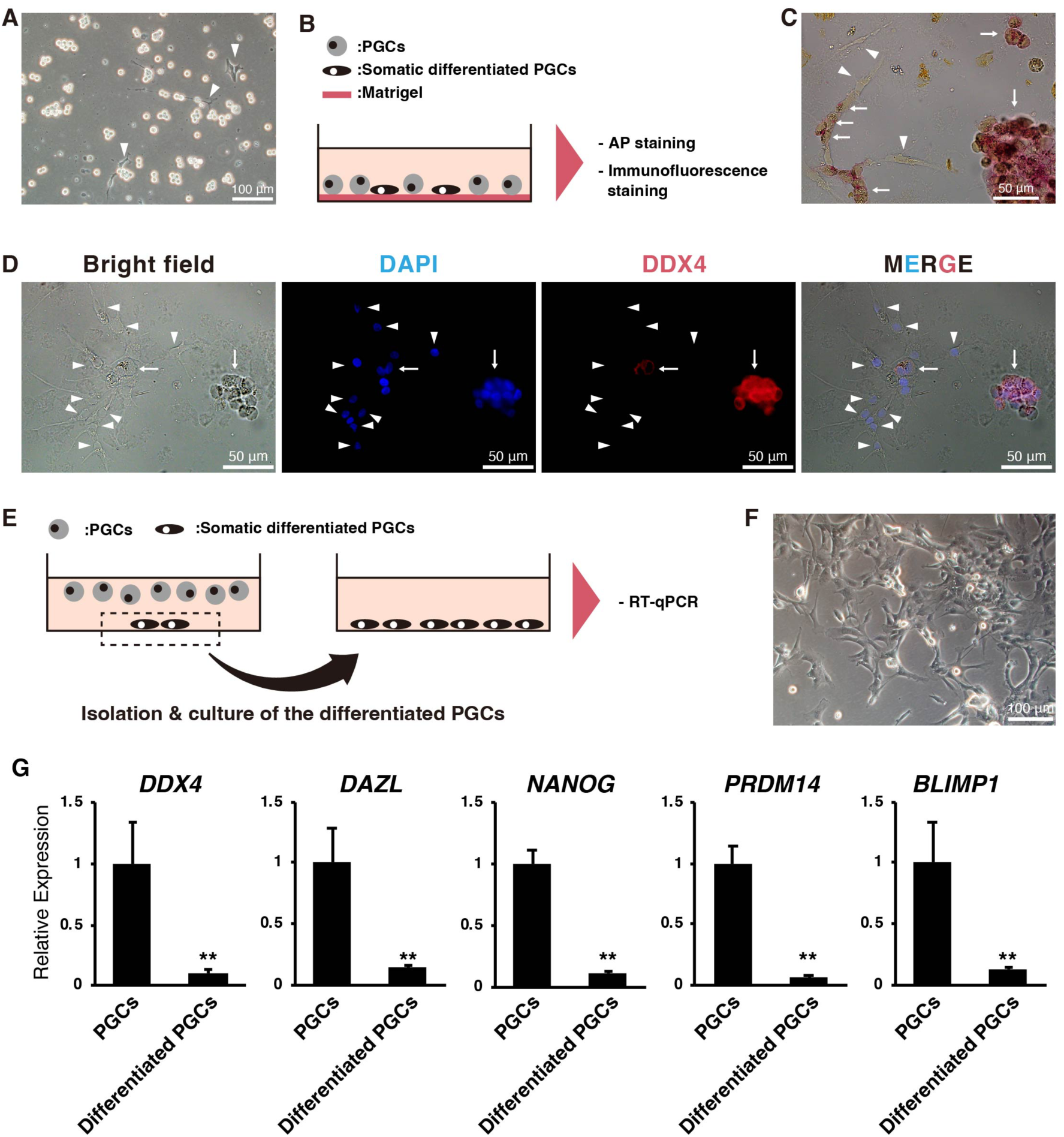
Characterization of somatic differentiated PGCs. (A) An image of somatic differentiated PGCs. The differentiated PGCs are indicated by arrowheads. (B) A schematic illustration of experimental design for the alkaline phosphatase and the immunofluorescence staining. The PGCs were seeded in an 8-well chamber slides with the Matrigel to adhere the cells into the bottom of the slides and incubated for 48 hours. The detail of experimental conditions is described in the Materials & Methods section. (C) Alkaline phosphatase staining of PGCs. Undifferentiated PGCs and the somatic differentiated PGCs are indicated by arrows and arrowheads, respectively. (D) Immunofluorescence staining of PGCs using anti-DDX4 antibody. Nuclear staining was performed with DAPI. Undifferentiated PGCs and the somatic differentiated PGCs are indicated by arrows and arrowheads, respectively. (E) A schematic illustration for isolation of the differentiated PGCs. As the somatic differentiated PGCs are adherent into a surface of dishes and undifferentiated PGCs typically float, somatic differentiated PGCs can be purified by removing medium supernatant with the floating cells. (F) An image of the isolated and proliferating differentiated PGCs. (G) Expression analysis of germ cell development-related genes. The ΔΔCt method was used to calculate relative expression levels. The levels were normalized to β-actin mRNA expression. Error bars indicate SD of the mean of expression levels in independently cultured PGCs derived from independent experiment (n = 3). Statistical significance was evaluated using the *t*-test. Asterisks represent significant differences (\*\**P* < 0.01).

We conducted a reverse transcription-quantitative polymerase chain reaction (RT-qPCR) analysis on these two populations of cells. To accomplish this, the adherent PGCs were enriched by removal of the suspension PGCs by aspiration (Fig. 2E) and continued proliferation for an additional two–three weeks (Fig. 2F). The RT-qPCR analysis revealed that the expression of other germ cell markers [23], *deleted in azoospermia like* (*DAZL*), *Nanog homeobox* (*NANOG*), *PR-domain-containing protein 14* (*PRDM14*), and *B lymphocyte-induced maturation protein 1* (*BLIMP1*), was also lost. (Fig. 2G). As expected, these results indicate that PGCs downregulate germ cell-specific genes as they acquire a somatic cell-like characters.

### 2.3. Gene expression profiling of the somatic differentiated chicken PGCs using RNA-seq

To identify the genetic pathways activated in PGCs undergoing somatic cell conversion, we conducted gene expression profiling using RNA-seq analysis on these two populations of cells. Approximately 45.9 million total reads per sample were obtained, and nearly 89.4% of these reads were uniquely mapped to the GRCg6a reference genome (Supplementary Table S1). A Pearson correlation coefficient of gene expression patterns between each sample was calculated (Fig. 3A), and the correlation was visualized by clustering analysis (Fig. 3B). As a result, each biological replicate was highly correlated (Pearson’s r ≥ 0.96) and clustered, indicating a reliable consistency of the sequencing.

**Fig. 3.**
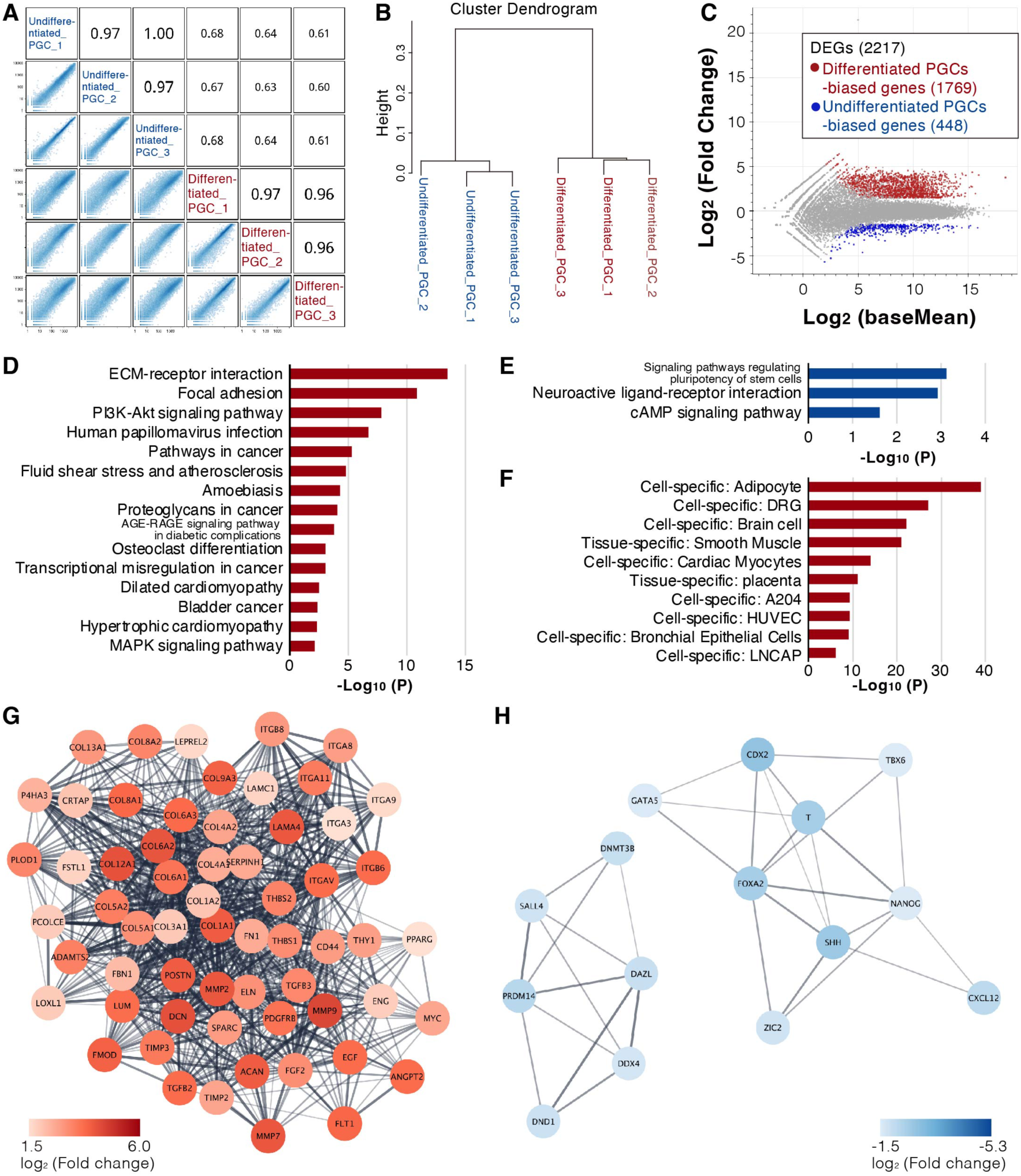
Gene expression profiling using RNA-seq. (A) Correlation diagram for each sample. (B) Hierarchical clustering of each sample based on gene expression profiles obtained by the RNA-seq analysis. (C) MA-plot of the somatic differentiated PGCs versus undifferentiated PGCs. The somatic differentiated PGCs-biased genes (log2 (FC) ≥ 1.5, Padj < 0.01) are shown in red. Undifferentiated PGCs-biased genes (log2(FC) ≤ −1.5, Padj < 0.01) are shown in blue. (D, E) The result of KEGG pathway analysis using the somatic differentiated PGCs-biased genes (D) and the undifferentiated PGCs-biased genes (E). (F) The result of tissue- or cell-specificity. (G, H) The protein–protein interaction network of the somatic differentiated PGCs-biased proteins (G) and the undifferentiated PGCs-biased proteins (H). Nodes and edges represent protein and interaction, respectively.

A total of 2217 differentially expressed genes (DEGs) (|log2 (Fold change)| ≥ 1.5, adjusted p-value (Padj) < 0.01) were identified, and 1769 and 448 genes of those showed the differentiated PGCs-biased and undifferentiated PGCs-biased expression, respectively (Fig. 3C). All DEGs detected in this study are listed in supplementary Table S2.

The identified DEGs were used for subsequent enrichment analyses. A Kyoto Encyclopedia of Genes and Genomes (KEGG) pathway analysis [24] was conducted using the differentiated PGCs-biased genes. The KEGG pathway analysis showed that cell-extracellular matrix (ECM) interaction-related pathways and cancer-related pathways, including the phosphatidylinositol-3-kinase (PI3K)-Akt signaling [25], were significantly enriched in the somatic differentiated PGCs (Fig. 3D). A gene ontology (GO) analysis using the same DEGs similarly showed significant enrichment of ECM related genes (Supplementary Fig. S2). In contrast, KEGG pathway analysis of the undifferentiated PGCs-biased genes showed an enrichment of stem cell-related pathways (Fig. 3E). Next, we analyzed cell- and tissue-specificity using the somatic differentiated PGCs-biased genes. Interestingly, the differentiated PGCs-biased genes were significantly enriched for several mesenchymal cell linages, such as adipocytes, smooth muscle, and cardiac monocytes, and also for neuron cells (Fig. 3F), further emphasizing that the differentiated PGCs were converted from a germ cell linage to a somatic linage. All genes enriched in each pathway, gene ontology, and cell- or tissue-specificity are listed in supplementary Table S3.

We next performed protein-protein interaction (PPI) network analyses using the undifferentiated- and somatic differentiated PGCs-biased genes. In the differentiated PGCs, the cluster of the PPI network was composed of integrins (ITGs), collagens (COLs), matrix metalloproteinases (MMPs), and several growth factors such as transforming growth factor betas (TGF-βs) (Fig. 3G). On the other hand, the clusters in the PPI network derived from the undifferentiated PGCs biased genes showed an enrichment of germ cell characteristic factors, including DNA methyltransferase 3 beta (DNMT3B), involved in *de novo* DNA methylation of chicken PGCs [26], the C-X-C motif chemokine ligand 12 (CXCL12), a factor involved in gonadal migration of germ cells [27, 28], in addition to the germ cell markers DDX4, DAZL, PRDM14, and NANOG (Fig. 3H).

### 2.4. Evaluation of partial EMT features of cultured PGCs

Our previous study showed that chicken PGCs express both E-cadherin, an epithelial cell marker, and N-cadherin, a mesenchymal cell marker [2]. In addition, the enrichment analysis revealed that the several somatic differentiated PGCs-biased genes were significantly enriched in some mesenchymal cells (Fig. 3F). Moreover, the up-regulation of ITGs, MMPs, and ECM components, shown in the PPI network analysis (Fig. 3G), during the conversion of PGCs is a typical feature of the EMT [29].Thus, we hypothesized that the PGCs underwent a partial epithelial-to-mesenchymal transition (EMT) during their somatic conversion, namely, the transition from an intermediate state expressing both epithelial and mesenchymal properties to a mesenchymal state.

To confirm this hypothesis, we used the RNA-seq data and compared expression patterns of multiple epithelial markers, such as *cadherin 1* (*CDH1*, the E-cadherin coding gene), *occludin* (*OCLN*), and *claudin 1* (*CLDN1*), and mesenchymal markers, such as *vimentin* (*VIM*) and *S100 calcium binding protein A4* (*S100A4*), between undifferentiated and differentiated PGCs. Also, we evaluated genes of *snail family transcriptional repressor 1* (*SNAI1*), *SNAI2*, *zinc finger E-box binding homeobox 1* (*ZEB1*), and *twist family bHLH transcription factor 1* (*TWIST1*), major transcription factors inducing the EMT and detected as mesenchymal markers [30]. We observed the suppression of many epithelial markers and activation of the mesenchymal markers were observed in the somatic differentiated PGCs, (Fig. 4A).

**Fig. 4.**
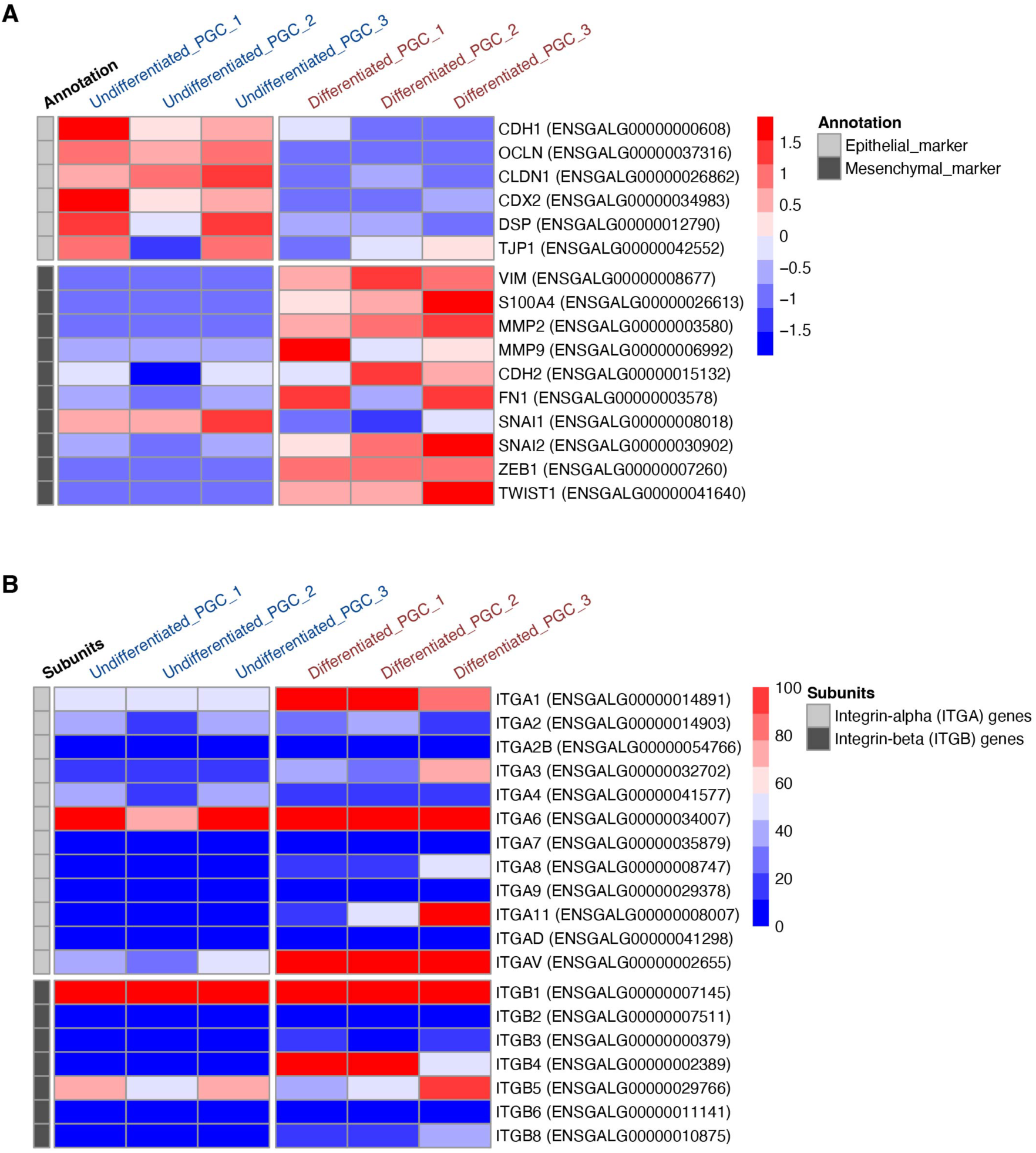
Comparison of expression patterns of EMT-related genes between undifferentiated and the somatic differentiated PGCs. (A) A heatmap of mesenchymal and epithelial markers for each sample. The red-blue color scale represents Z-score of transcripts per kilobase million (TPM) values, with red indicating high and blue indicating low expression levels. (B) A heatmap of integrin genes for each sample. The red-blue color scale represents TPM, with red indicating high and blue indicating low expression levels.

We further investigated the alteration of expression patterns of integrins, heterodimers of integrin alpha- and beta-subunits, during the somatic lineage conversion, as this is a defining characteristic of EMT progression [29]. An heatmap analysis showed upregulation of additional integrin genes, such as *integrin subunit alpha V* (*ITGAV*) and *integrin subunit beta 4* (*ITGB4*), during the conversion, suggesting that the somatic differentiated PGCs acquired additional integrin heterodimers (Fig. 4B). Interestingly, the undifferentiated PGCs, semi-adherent suspension cells in vitro, also exhibited the expression of several integrin genes, particularly *ITGA6* and *ITGB1*. Overall, these heatmap analyses supported our hypothesis that the somatic conversion of the chicken PGCs occurred through a partial EMT feature.

### 2.5. Confirmation of the RNA-seq results

To confirm the reliability of the RNA-seq data, we conducted RT-qPCR analyses. Here, we used three new independent chicken PGCs lines (n = 3). We targeted 16 DEGs, and the expression levels were compared between undifferentiated PGCs and the somatic differentiated PGCs. As a result, the expression patterns of all these DEGs in the RT-qPCR analysis corresponded to those in the RNA-seq analysis (Fig. 5A). Subsequently, we evaluated the protein expression of the mesenchymal marker, VIM, in the somatic differentiated PGCs using immunofluorescence analysis. We observed specific expression of VIM in the adherent PGCs (Fig. 5B), also suggesting the partial-EMT feature of the somatic conversion.

**Fig. 5.**
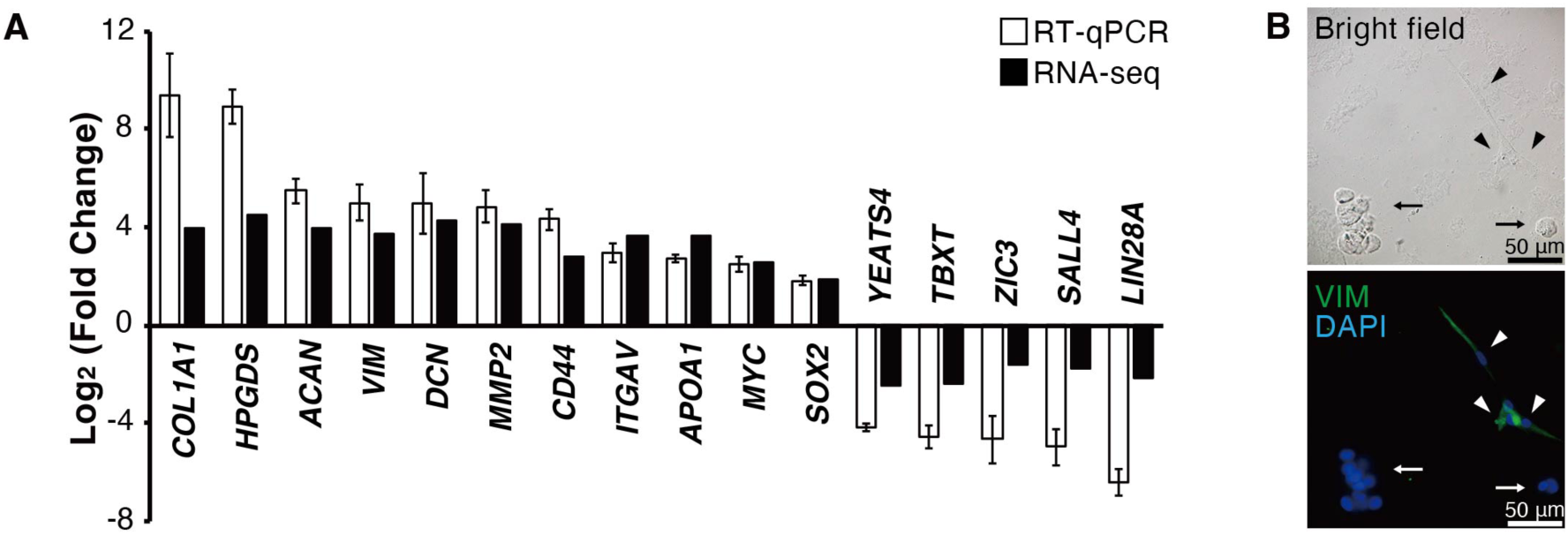
Expression analysis of DEGs. (A) RT-qPCR analysis targeting 16 DEGs. The y-axis indicates the log2 ratio of the somatic differentiated PGCs-to-undifferentiated PGCs expression levels for each gene. Open bars indicate the results of RT-qPCR, and closed bars indicates the result of RNA-seq. For the RT-qPCR analysis, the ΔΔCt method was used to calculate the expression levels. The levels were normalized to β-actin mRNA expression. Error bars indicate SD of the mean of expression levels in independently cultured PGCs derived from three embryos (n = 3). (B) Immunofluorescence staining of chicken PGCs using anti-VIM antibody. Counter staining was performed with DAPI. Undifferentiated PGCs and the somatic differentiated PGCs are indicated by arrows and arrowheads, respectively.

### 2.6. Identification of factors inducing the somatic differentiation in chicken PGCs

We next investigated matrix factors inducing the somatic conversion of chicken PGCs. Previous studies demonstrated that a cell-extracellular matrix (ECM) interaction and its associated mechanical stress induced EMT [31, 32]. Thus, we hypothesized that the interaction of chicken PGCs with the surface of the dish causes somatic conversion through a partial EMT.

For this experiment we used *PRDM14^eGFP/+^* chicken PGCs [33]. The *PRDM14^eGFP/+^* PGCs express green fluorescence in their undifferentiated state, whilst the expression is quickly lost after somatic differentiation (Fig. 6A). PGCs were cultured in proliferation medium on fibronectin (FN), an ECM component, to induce adhesion of the PGCs on the surface of the culture dish. As a result, microscopic observation showed that the PGCs cultured with FN frequently underwent the somatic differentiation in a time-dependent manner (Fig. 6B). The flow cytometric (FCM) analysis also revealed a significant increase of eGFP-negative PGCs, namely somatic converted PGCs, under the FN coat condition (Fig. 6C & D). The same tendency was also observed in the PGCs cultured with collagen type I (Supplementary Fig. S3). These results indicated that one factor driving the somatic differentiation of PGCs is an interaction of the cell membrane with the matrix. Also, given that inhibiting a PGC-matrix interaction reduced their self-renewal capacity (Fig. 1), this suggests that whilst the interaction is an essential factor for PGC proliferation in vitro, a robust PGC-matrix interaction triggers disruption of the PGC self-renewal pathway (Fig. 7).

**Fig. 6.**
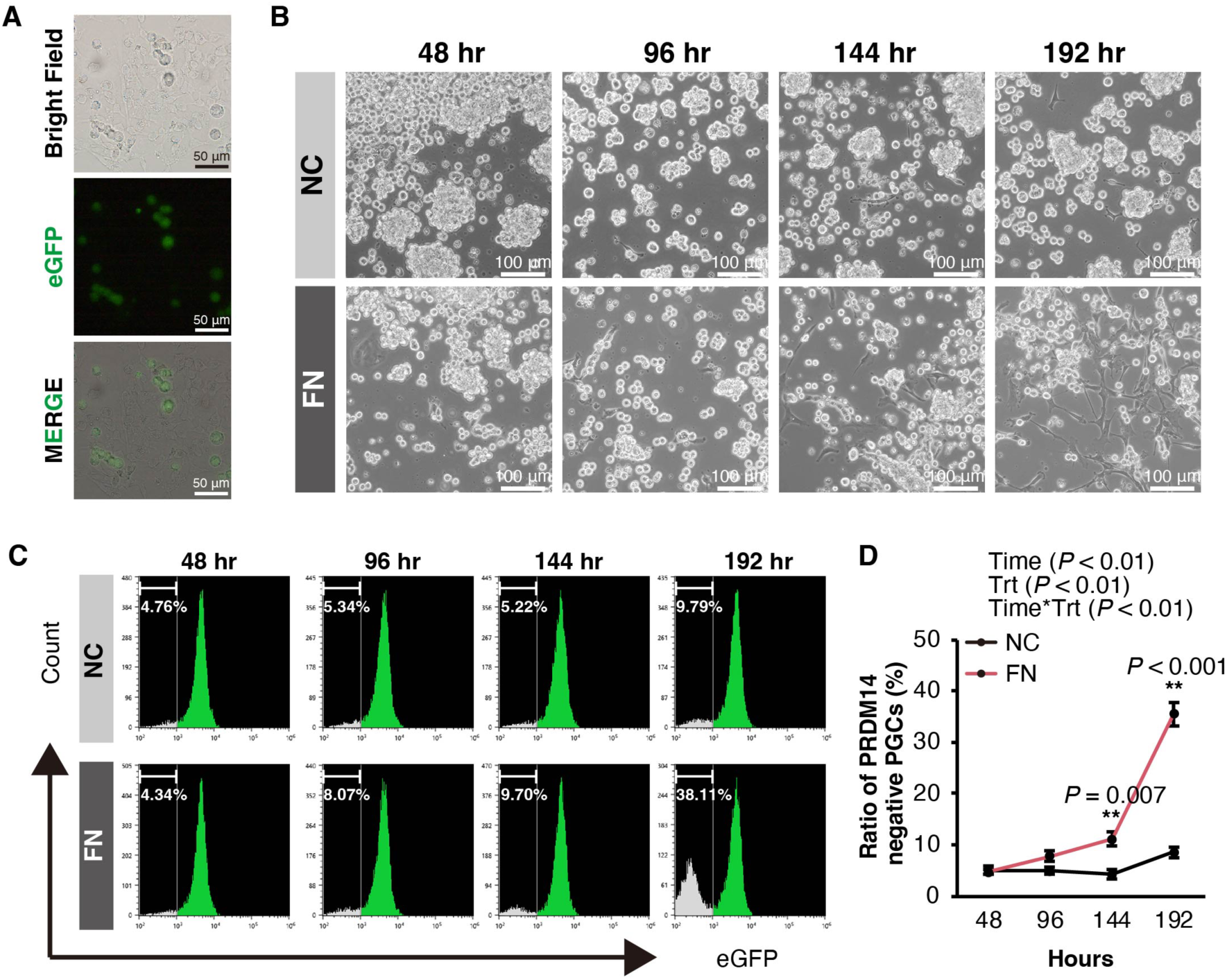
Analyses of a PGC-matrix interaction. (A) Fluorescence scoping of the cultured *PRDM14^eGFP^*^/+^ PGCs. (B) Photographs of the *PRDM14^eGFP^*^/+^ PGCs cultured with or without fibronectin in each time point. NC, negative control, and FN, fibronectin coating. (c, d) Results of flow cytometric analysis. Histograms (C) and a graph (D) show ratio of PRDM14 negative PGCs in each treatment and time point. Error bars indicate SD of the mean of the ratio of PRDM14 negative cells of three replicates (n = 3). The result of a two-way repeated measures ANOVA is shown at the top of the graph. Statistical significance in each time point was evaluated using Bonferroni’s post-hoc test. Asterisks represent significant differences (\*\**P* < 0.01). Trt, treatment.

**Fig. 7.**
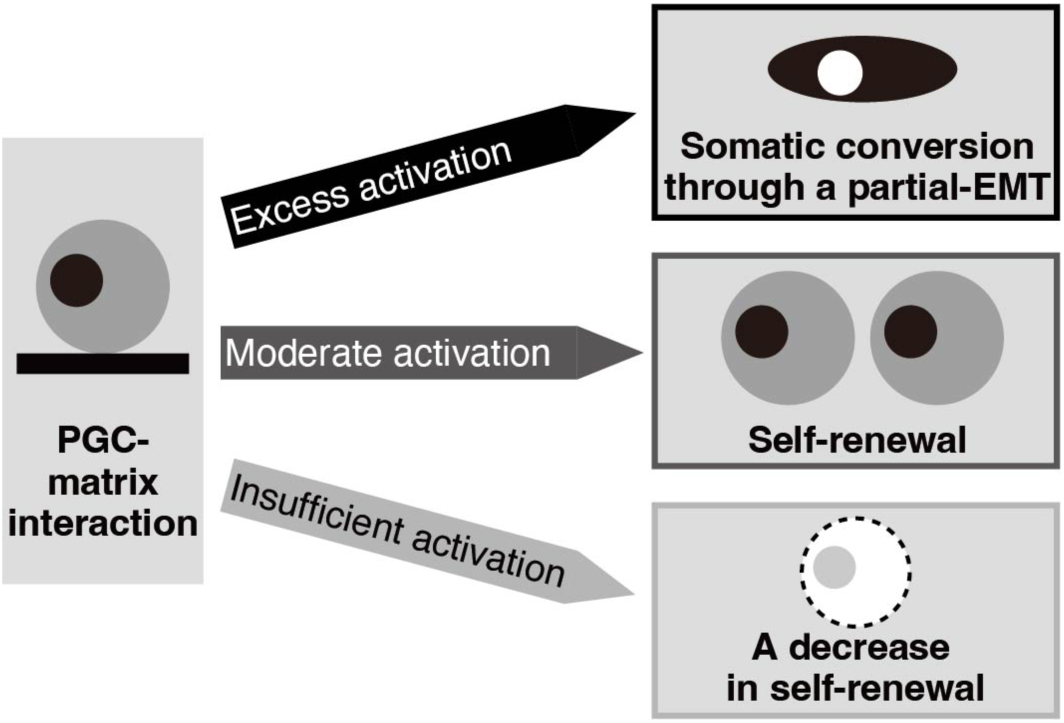
A model for the fate decision of avian primordial germ cells through their interaction with the matrix. For the stable culture of avian PGCs, an appropriate level of cell-matrix interaction is required; otherwise, somatic conversion or reduction in the self-renewal capacity occurs.

## 3. Discussion

Establishing a culture system for PGCs from other avian species will aid to develop other avian model systems and aiding the conservation of avian genetic resources [10]. However, due to a poor self-renewal capacity and the instability of maintaining undifferentiated states, propagation of non-chicken avian PGCs remains unachieved [11, 12]. Thus, we investigated the fate decision process of chicken PGCs and revealed that a cell-matrix interaction is a factor regulating the self-renewal and somatic conversion of avian PGCs (Fig. 7). Our results provide valuable insights for improving a culture system for avian PGCs.

### 3.1. Conversion of avian PGCs to somatic cell lineages through a partial EMT

A characterization of germ cell markers showed that their expression was lost in the somatic-differentiated PGCs (Fig. 2), indicating that the adherent PGCs which have fibroblast-like morphology lacked germ-cell properties. Further RNA-seq analysis suggested that the cultured PGCs initiated the somatic conversion through a partial EMT (Fig. 3, 4 & 5). EMT is shown to govern several cell type conversions, such as cancer cells during metastasis, epiblast cells during gastrulation, and pluripotent stem cells during their induction. Typical properties of these cells during EMT are the disassembly of cell-cell junctions and acquisition of cellular motility, accompanied by down-regulation of epithelial markers and up-regulation of mesenchymal markers [29]. Although the chicken PGCs are not epithelial cells, they express both E-cadherin, an epithelial marker, and N-cadherin, a mesenchymal marker [2]. Thus, we referred to this phenomenon as a partial EMT: a transition from intermediate states between epithelial and mesenchymal into a mesenchymal state. Note that an EMT can exhibit various degrees, from full EMT to partial EMT, and a partial EMT is the standard status rather than the exception in cancer cells [34].

In the PPI network constructed by the somatic differentiated PGCs-biased proteins, ITGs, ECM components, MMPs, and several growth factors were significantly enriched (Fig. 3G). These molecules are also upregulated during typical EMT, and some of them contribute to its progression. Remodeling the properties of a cell-ECM interaction, namely an integrin-ECM interaction, is an essential step for initial EMT [29]. The activation of collagens and fibronectins are defined as biomarkers of EMT [35]. The expression of MMPs is activated during EMT, enhancing the movement and invasion of cells through proteolysis of ECM proteins [36]. Furthermore, the growth factors, particularly the TGF-β family [37–39], are known as EMT inducers, and the TGF-βs can progress EMT in the paracrine and autocrine manner [40]. Thus, the enrichment of these factors in the somatic differentiated PGCs reflects the existence of a partial EMT feature during the conversion.

### 3.2. A fate decision process in avian PGCs through a cell-matrix interaction

Inhibition of a PGC-matrix interaction and FAK signaling, a major pathway under integrin activation, reduced a self-renewal capacity (Fig. 1). In addition, our RNA-seq data show that the undifferentiated PGCs possess α6β1 integrin (Fig. 4B) that has a fundamental role for self-renewal in mammalian pluripotent stem cells [41, 42]. These results indicated that a cell-matrix interaction has a pivotal role for their in vitro propagation. In contrast, culturing the PGCs on matrix-coated plates in physiological normal Ca^2+^ condition caused an increase in somatic conversion (Fig. 6). A recent study showed that quail PGCs express several integrin subunits and matrix components and modulate ECM environment in early embryonic stages [14]. Also, tenascin-C, a glycoprotein acting as a negative regulator of cell-matrix interactions, is localized along the migratory route of chicken PGCs, and its role in modulating the motility of the PGCs has been suggested [43]. These insights shown in avian embryos imply that regulation of the signaling in a cell-matrix interaction is vital for PGC development, which is compatible with our current findings. Our findings also suggested the existence of the ‘Goldilocks principle’ in the context of the strength of signaling in a cell-matrix interaction for the stable growth of avian PGCs, and thus regulating the interaction and its downstream signals may become a key for the culture of non-chicken avian PGCs.

In this study, we revealed that a self-renewal capacity and undifferentiated states of avian PGCs are regulated by a PGC-ECM interaction. However, the signaling pathway(s) and master regulatory factor(s) directly regulating this fate decision remain unclear although FAK phosphorylation is needed. Also, a recent gene expression profiling exhibited different characteristics of PGCs across avian species [44], suggesting that the strength of the signaling pathway(s) for stable self-renewal of PGCs needs to be optimized in each species. Thus, further molecular studies will be needed to understand the repression of the somatic conversion of avian PGCs without reduction of PGC self-renewal potentially through chemical inhibitors.

## 4. Materials & Methods

### 4.1. Experimental animals

Fertilized eggs of chickens (White-Leghorn) were purchased from Akita foods (Fukuyama, Japan) or National Avian Research Facility (NARF) at the Roslin Institute (Scotland). The eggs of the *PRDM14^eGFP/+^* genome-edited chickens were purchased from the NAGOYA UNIVERSITY through the National Bio-Resource Project of the MEXT, Japan. All Japan animal experiments in this study were conducted in accordance with the guidelines of the Hiroshima University Animal Research Committee (approved protocol ID: C23-38). All UK animal management, maintenance and embryo manipulations were carried out under UK Home Office license PP9565661 and regulations. Experimental protocols and studies were approved by the Roslin Institute Animal Welfare and Ethical Review Board Committee.

### 4.2. Cell culture

Chicken PGCs were isolated from blood of chicken embryos at Hamburger and Hamilton stage (HH) 16 to 16^+^. In the high calcium condition (1.8 mM), the PGCs were maintained in a culture media established by Whyte et al. [2] with some modification described in our previous study [19]. Briefly, the KnockOut DMEM (Thermo Fisher Scientific Carlsbad, CA, USA) was supplemented with 1% chicken serum (Thermo Fisher Scientific), 1 × B-27 (Thermo Fisher Scientific), 2 mM GlutaMAX™ (Thermo Fisher Scientific), 1 × nucleosides (Merck, Darmstadt, Germany), 1 × MEM Non-Essential Amino Acids Solution (NEAA) (Thermo Fisher Scientific), 1 × antibiotic–antimycotic mixed stock solution (Nacalai Tesque, Kyoto, Japan), 0.5 mM monothioglycerol (FUJIFILM Wako Pure Chemical Co., Osaka, Japan), 1 mM sodium pyruvate, 1 unit/mL heparin (Merck), 10 ng/mL human FGF2 (PeproTech Inc., Rocky Hill, NJ, USA), 0.2 μM H1152 (FUJIFILM Wako Pure Chemical Co.), and 0.2 μM Blebbistatin (FUJIFILM Wako Pure Chemical Co.). The adherent PGCs were also maintained in this condition and maintained for 2-3 weeks after isolation by aspiration of suspension PGCs and then they were used for subsequent RT-qPCR and RNA-seq analyses. In low calcium conditions (0.15 mM), the PGCs were cultured in the FAOT (FGF2, Activin, and Ovotransferrin) medium using the no-calcium containing avian knockout DMEM with 0.15 mM CaCl_2_, which is established by Whyte et al. [2]. The PGCs were cultured at 38°C with 5% CO_2_ and subcultured every 2-3 days. Cultured PGCs were observed and photographed using an Olympus IX71 inverted microscope with an Olympus DP70 camera (Olympus, Tokyo, Japan) and a Zeiss Axiovert 5 inverted microscope with a Zeiss Axiocam 395 mono camera (Zeiss, Oberkochen, Germany).

### 4.3. Immunofluorescence staining and AP staining

The anti-DDX4 [45] and anti-VIM (sc-80975; Santa Cruz Biotechnology, Dallas, TX, USA) monoclonal antibodies were used for the detection of each protein. First, cultured chicken PGCs were seeded into Millicell® EZ SLIDES (Merck) at a density of 1.0 × 10^4^ cells/well in the culture media with 1% Matrigel (Corning Inc., Corning, NY, USA) to adhere the cells to the surface of the slide, and the cells were cultured for 48 hr. Subsequently, cells were washed with PBS and fixed with 4% paraformaldehyde in PBS for 30 min at room temperature (approximately 20°C). After washing with 10 mM glycine-PBS, cells were incubated with 0.1% Triton X for 5 min at room temperature. Subsequently, the cells were washed with PBS and incubated with 3% BSA-PBS for 15 min at room temperature. Then, primary antibody incubation using hybridoma supernatant containing the anti-DDX4 antibody, which was diluted 1:1 with 2% BSA-PBS, or the anti-VIM antibody diluted 1:50 with 1% BSA-PBS was conducted for 40 min at 37°C. After washing, the secondary antibody incubation was performed using goat anti-mouse IgG (Alexa Fluor Plus 555; Invitrogen, Waltham, MA, USA) or goat anti-mouse IgM (Alexa Fluor 488; Invitrogen) at 1:200 dilution with 1% BSA-PBS for 40 min at 37°C. Finally, after washing, counterstaining was performed using DAPI in VECTASHIELD Mounting Medium (Vector Laboratories Inc., Burlingame, CA, USA).

The AP staining was performed using the Stemgent™ Alkaline Phosphatase staining kit II (ReproCELL, Yokohama, Japan). Cultured PGCs were seeded and cultured into Millicell^®^ EZ SLIDES (Merck) in the same condition described above, and the staining was performed according to the manufacturer’s instructions. Cells were observed under a microscope BX53 (Olympus) and imaged with a DP74 camera (Olympus).

### 4.4. RT-qPCR analysis

Gene expression analysis of the germ cell-development related genes and the DEGs detected in the RNA-seq analysis was conducted using RT-qPCR. Total RNA was purified using FastGene™ RNA Premium kit (NIPPON Genetics Co. Ltd., Tokyo, Japan). 100– 120 ng of purified RNA was used for cDNA synthesis using SuperScript IV reverse transcriptase (Thermo Fisher Scientific). RT-qPCR was performed using the StepOne™ real-time PCR system (Applied Biosystems Waltham, MA, USA) with the KOD SYBR^®^ qPCR Mix (Toyobo Co. Ltd. Osaka, Japan). The primers used in this analysis are listed in supplementary Table S4. The RT-qPCR reaction was performed under the following conditions: 40 cycles of 98°C for 10 sec, 60–65 °C for 10 sec, and 68°C for 30 sec, followed by melt curve analysis in three steps, 95 °C for 15 sec, 60 °C for 1 min, and 95 °C for 15 sec. The expression levels of each gene were normalized to those of *β-actin* and calculated using the ΔΔCt method [46].

### 4.5. RNA-seq and bioinformatic analysis

For the RNA-seq analysis, cultured PGCs derived from three individual embryos were used (biological replicates = 3), and the somatic differentiated PGCs were isolated from each of the PGC-lines, respectively. 1 μg of total RNA was purified using FastGene™ RNA Premium kit (NIPPON Genetics Co. Ltd., Tokyo, Japan) from each PGC line, after culture for two months. Subsequently, mRNA was enriched from the total RNA using NEBNext^®^ Poly(A) mRNA Magnetic Isolation Module (New England Biolabs, Ipswich, MA, USA), and cDNA libraries were constructed using SMARTer^®^ Stranded RNA-Seq Kit (TaKaRa Bio, Kusatsu, Japan) and MGIEasy Universal Library Conversion Kit (MGI, Shenzhen, China). These were analyzed using the DNBSEQ (MGI) with 150 bp paired-end sequencing.

The quality of the row sequence reads was assessed using FastQC (ver. 0.11.9). The raw reads were trimmed and quality filtered using Trimmomatic (ver. 0.39) [47] as well as Trim Galore (ver. 0.6.6) with Cutadapt (ver. 3.1) [48] software then mapped to the chicken genome (GRCg6a) using STAR (ver. 2.7.3a) [49]. Reads were counted using RSEM (ver.1.3.1) [50].

Detection of DEGs were performed using the DESeq2 tool (ver. 1.20.0) [51]. MA-plot was created using the Bokeh visualization library (ver. 2.4.2). Clustering and correlationship analysis were performed using R software (https://www.r-project.org) with the TCC package (ver. 1.14.0) [52] based on read counts normalized by the DESeq2 tool. In this analysis, total 6424 of significantly variable genes across samples (Padj < 0.05) detected using the DESeq2 tool was used. KEGG pathway and GO analyses were performed using the g:Profiler online tool [53], and tissue or cell-specificity was characterized using the Metascape online tool [54]. PPI network analysis was performed using MCODE (Degree Cutoff: 2, Node Score Cutoff: 0.3, K-Core: 2, Max. Depth: 100) through Cytoscape software (ver 3.9.0) based on the STRING database (https://string-db.org). The heatmap was created in R software using the pheatmap package (ver 1.0.12) [55].

### 4.6. ECM reagents and flowcytometric analysis

Human plasma fibronectin (Merck) and bovine skin collagen type I (Nippi Co. Ltd., Tokyo, Japan) were used to coat the surface of the culture dishes. The fibronectin was diluted by PBS and coated in culture dishes with 5 μg/cm^2^. On the other hand, the collagen was diluted with 5 mM acetic acid at a dose of 0.01%, and 1 mL of the collagen solution was applied to 12-well plates. For coating, they were incubated overnight at room temperature (approximately 20°C).

In the culture of PGCs with the matrix reagents, PGCs derived from *PRDM14^eGFP/+^*genome-edited chickens and cultured for about two months were used. The PGCs were seeded into a 12-well plate with or without the ECM coating at a density of 1.0 × 10^4^ cells/well. The PGCs cultured in each condition (n = 3) were collected every 48 hours. Half of the collected PGCs from each well were subcultured, and the other half of them were used for the flow cytometric analysis. The PGCs were washed with PBS containing 0.5% BSA and 0.1% NaN3, and the ratio of eGFP-negative PGCs was determined using MA900 Multi-Application Cell Sorter (Sony Biotechnology, San Jose, CA, USA).

### 4.7. Culturing PGCs with agarose gel and inhibitors

The 2% (w/v) agarose (Thermo Fisher Scientific) solution diluted by distilled water was applied into a 48-well tissue culture plate at a volume of 250 uL/well and incubated for 1 hr at room temperature. Wells were washed and equilibrated with the FAOT medium overnight.

Defactinib (ApexBio, Houston, TX, USA) was dissolved with DMSO at 1 mM, and the solution was added to the PGC culture medium at the density of 0.1% (v/v) (final concentration, 1 μM). The PGC culture medium including 0.1% DMSO was used as a negative control. In each of the experiments, the PGCs were seeded into a 48-well plate at a density of 2.0 × 10^2^ cells/well and maintained in the FAOT medium for 10 days. Then, the number of PGCs was counted by hand using a hemocytometer.

### 4.8. Western blotting

For the sample preparation, chicken PGCs were seeded into a 24-well plate at 1.5 × 10^5^ cells/well and cultured with the FAOT with 1 μM defactinib or 0.1% DMSO. After the incubation for 24 hrs, the number of PGCs in each sample was counted. Cells were then lysed in RIPA Lysis buffer (Santa Cruz Biotechnology) with 1 × Halt™ Protease and Phosphatase Inhibitor Cocktail (Thermo Fisher Scientific) at a concentration of 1.0 × 10^4^ cells/µL.

15 µL of each sample was mixed with beta-mercaptoethanol and NuPAGE™ LDS Sample Buffer (4X) (Thermo Fisher Scientific) and heated at 95°C for 5 min. After separating protein bands on a NuPAGE Bis-Tris Mini Protein Gel (4–12%) (Thermo Fisher Scientific), they were transferred onto a PVDF membrane (0.45 µm pore size) (Thermo Fisher Scientific). The membrane was subsequently blocked in 5% bovine serum albumin (BSA)/TBST at room temperature (approximately 20°C) for 1 hr. For detection of the phospho-FAK, the membrane was incubated in anti-phospho-FAK (Tyr397) polyclonal antibody (#44-624G) (Thermo Fisher Scientific) diluted at 1:1000 in 5% BSA/TBST overnight at 4°C. After washing with TBST, the membrane was incubated with anti-rabbit IgG, HRP-linked antibody (#7074) (Cell Signaling Technology, Danvers, MA, USA) diluted at 1:2000 in 5% BSA/TBST at room temperature for 1 hr. Following washing, the membrane was incubated with SignalFire™ ECL Reagent (Cell Signaling Technology) to initiate chemiluminescence. The chemiluminescence was detected using Amersham Hyperfilm™ ECL (GE Healthcare, Chicago, IL, USA) and an Optimax 2010 automated X-ray film processor (Protec, Oberstenfeld, Germany).

After the detection of the protein bands, the membrane was rinsed and stripped with Restore™ PLUS Western blot stripping buffer (Thermo Fisher Scientific), the membrane was incubated in anti-total-FAK polyclonal antibody (#12636-1-AP) (Proteintech, Rosemont, IL, USA) diluted at 1:100000 in 5% BSA/TBST overnight at 4°C. A secondary antibody reaction and detection of the chemiluminescence were conducted using the same procedure described above.

### 4.9. Statistical analysis

Statistical analyses were performed using EZR software (ver 1.61) [56]. Data obtained by RT-qPCR analysis were analyzed using the *t*-test. Data obtained by FCM analysis were analyzed using the two-way repeated measures analysis of variance (ANOVA) with respect to time (hours of culture) and treatments (with or without ECM reagents). followed by the Bonferroni post-test at each time point. A *p*-value < 0.05 was considered significant.

## Acknowledgement

K.I. was funded by the Japan Society for the Promotion of Science (JSPS) Overseas Research Fellowship and the Kieikai Research Foundation (FY2022). H.H. was funded by JSPS KAKENHI under grant 19H03107, NBRP (chicken & Quail) and JST COI under grant JPMJPF 2010. M.J.M was funded by Institute Strategic Grant Funding from the BBSRC (BB/P0.13732/1 and BB/P013759/1) to the Roslin Institute. Project funding was also obtained from Colossal Biosciences (USA). We would like to thank K.K. DNAFORM (Yokohama, Japan) for the RNA-seq analyses.

## CRediT authorship contribution statement

**Kennosuke Ichikawa:** Conceptualization, Methodology, Validation, Formal analysis, Investigation, Resources, Data Curation, Writing - Original Draft, Writing - Review & Editing, Visualization, Project administration, Funding acquisition. **Yuzuha Motoe:** Methodology, Validation, Formal analysis, Investigation, Resources. **Tenkai Watanabe**: Methodology, Resources. **Ryo Ezaki**: Methodology, Resources. **Mei Matsuzaki**: Methodology, Resources. **Hiroyuki Horiuchi:** Conceptualization, Methodology, Supervision, Funding acquisition. **Michael J. McGrew:** Conceptualization, Methodology, Resources, Writing - Review & Editing, Supervision, Funding acquisition

## Data availability

All RNA-seq datasets are publicly available at Gene Expression Omnibus under accession number GSE280091.

## References

1. van de Lavoir MC, Diamond JH, Leighton PA, Mather-Love C, Heyer BS, Bradshaw R, Kerchner A, Hooi LT, Gessaro TM, Swanberg SE et al: Germline transmission of genetically modified primordial germ cells. Nature 2006, 441(7094):766-769. 10.1038/nature04831

2. Whyte J, Glover JD, Woodcock M, Brzeszczynska J, Taylor L, Sherman A, Kaiser P, McGrew MJ: FGF, Insulin, and SMAD Signaling Cooperate for Avian Primordial Germ Cell Self-Renewal. Stem Cell Rep 2015, 5(6):1171–1182. 10.1016/j.stemcr.2015.10.008

3. Ichikawa K, Matsuzaki M, Ezaki R, Horiuchi H: Genome editing in chickens. Gene and Genome Editing 2022, 3–4:100015. 10.1016/j.ggedit.2022.100015

4. Woodcock ME, Gheyas AA, Mason AS, Nandi S, Taylor L, Sherman A, Smith J, Burt DW, Hawken R, McGrew MJ: Reviving rare chicken breeds using genetically engineered sterility in surrogate host birds. Proc Natl Acad Sci U S A 2019, 116(42):20930–20937. 10.1073/pnas.1906316116

5. Webster RG, Bean WJ, Gorman OT, Chambers TM, Kawaoka Y: Evolution and ecology of influenza A viruses. Microbiol Rev 1992, 56(1):152–179. 10.1128/mr.56.1.152-179.1992

6. Barber MR, Aldridge JR, Jr., Webster RG, Magor KE: Association of RIG-I with innate immunity of ducks to influenza. Proc Natl Acad Sci U S A 2010, 107(13):5913–5918. 10.1073/pnas.1001755107

7. Ichikawa K, Motoe Y, Ezaki R, Matsuzaki M, Horiuchi H: Knock-in of the duck retinoic acid-inducible gene I (RIG-I) into the Mx gene in DF-1 cells enables both stable and immune response-dependent RIG-I expression. Biochem Biophys Rep 2021, 27:101084. 10.1016/j.bbrep.2021.101084

8. Woo SJ, Choi HJ, Park YH, Rengaraj D, Kim J-K, Han JY: Amplification of immunity by engineering chicken MDA5 combined with the C terminal domain (CTD) of RIG-I. Appl Microbiol Biotechnol 2022, 106(4):1599–1613. 10.1007/s00253-022-11806-4

9. Okuno M, Miyamoto S, Itoh T, Seki M, Suzuki Y, Mizushima S, Kuroiwa A: Expression profiling of sexually dimorphic genes in the Japanese quail, Coturnix japonica. Sci Rep 2020, 10(1):20073. 10.1038/s41598-020-77094-y

10. Ichikawa K, McGrew MJ: Innovations in poultry reproduction using cryopreserved avian germ cells. Reprod Domest Anim 2024, 59(5), e14591. 10.1111/rda.14591

11. Chen Y-C, Lin S-P, Chang Y-Y, Chang W-P, Wei L-Y, Liu H-C, Huang J-F, Pain B, Wu S-C: In vitro culture and characterization of duck primordial germ cells. Poult Sci 2019, 98(4):1820–1832. 10.3382/ps/pey515

12. Park TS, Kim MA, Lim JM, Han JY: Production of quail (Coturnix japonica) germline chimeras derived from in vitro-cultured gonadal primordial germ cells. Mol Reprod Dev 2008, 75(2):274–281. 10.1002/mrd.20821

13. Naito M, Harumi T, Kuwana T: Long Term in vitro Culture of Chicken Primordial Germ Cells Isolated from Embryonic Blood and Incorporation into Germline of Recipient Embryo. J Poult Sci 2010, 47(1):57–64. 10.2141/jpsa.009058

14. Huss DJ, Saias S, Hamamah S, Singh JM, Wang J, Dave M, Kim J, Eberwine J, Lansford R: Avian primordial germ cells contribute to and interact with the extracellular matrix during early migration. Front Cell Dev Biol 2019, 7: 35. 10.3389/fcell.2019.00035

15. Choi JW, Kim S, Kim TM, Kim YM, Seo HW, Park TS, Jeong J-W, Song G, Han JY: Basic Fibroblast Growth Factor Activates MEK/ERK Cell Signaling Pathway and Stimulates the Proliferation of Chicken Primordial Germ Cells. PLoS One 2010 5(9): e12968. 10.1371/journal.pone.0012968

16. Macdonald J, Glover JD, Taylor L, Sang HM, McGrew MJ: Characterisation and Germline Transmission of Cultured Avian Primordial Germ Cells. PLoS One 2010 5(11): e15518. 10.1371/journal.pone.0015518

17. Costa EC, de Melo-Diogo D, Moreira AF, Carvalho MP, Correia IJ: Spheroids Formation on Non-Adhesive Surfaces by Liquid Overlay Technique: Considerations and Practical Approaches. Biotechnol J 2018 13(1): 1700417. 10.1002/biot.201700417

18. Vitillo L, Kimber SJ: Integrin and FAK Regulation of Human Pluripotent Stem Cells. Curr Stem Cell Rep 2017 3(4): 358–365. 10.1007/s40778-017-0100-x

19. Ezaki R, Hirose F, Furusawa S, Horiuchi H: An improved protocol for stable and efficient culturing of chicken primordial germ cells using small-molecule inhibitors. Cytotechnology 2020, 72(3):397–405. 10.1007/s10616-020-00385-9

20. Tang X, Zhang C: Activation of protein kinases A and C promoted proliferation of chicken primordial germ cells. Anim Reprod Sci 2007, 101(3):295–303. 10.1016/j.anireprosci.2006.09.020

21. Yu F, Zhu Z, Chen X, Huang J, Jia R, Pan J: Isolation, characterization and germline chimera preparation of primordial germ cells from the Chinese Meiling chicken. Poult Sci 2019, 98(2):566–572. 10.3382/ps/pey410

22. Tsunekawa N, Naito M, Sakai Y, Nishida T, Noce T: Isolation of chicken vasa homolog gene and tracing the origin of primordial germ cells. Development 2000, 127(12):2741–2750. 10.1242/dev.127.12.2741

23. Ichikawa K, Horiuchi H: Fate Decisions of Chicken Primordial Germ Cells (PGCs): Development, Integrity, Sex Determination, and Self-Renewal Mechanisms. Genes 2023, 14(3):612. 10.3390/genes14030612

24. Kanehisa M, Goto S: KEGG: Kyoto Encyclopedia of Genes and Genomes. Nucleic Acids Res 2000, 28(1):27–30. 10.1093/nar/28.1.27

25. Xu W, Yang Z, Lu N: A new role for the PI3K/Akt signaling pathway in the epithelial-mesenchymal transition. Cell Adh Migr 2015, 9(4):317–324. 10.1080/19336918.2015.1016686

26. Rengaraj D, Lee BR, Lee SI, Seo HW, Han JY: Expression patterns and miRNA regulation of DNA methyltransferases in chicken primordial germ cells. PLoS One 2011, 6(5):e19524. 10.1371/journal.pone.0019524

27. Stebler J, Spieler D, Slanchev K, Molyneaux KA, Richter U, Cojocaru V, Tarabykin V, Wylie C, Kessel M, Raz E: Primordial germ cell migration in the chick and mouse embryo: the role of the chemokine SDF-1/CXCL12. Dev Biol 2004, 272(2):351–361. 10.1016/j.ydbio.2004.05.009

28. Lee JH, Park JW, Kim SW, Park J, Park TS: C-X-C chemokine receptor type 4 (CXCR4) is a key receptor for chicken primordial germ cell migration. J Reprod Dev 2017, 63(6):555–562. 10.1262/jrd.2017-067

29. Lamouille S, Xu J, Derynck R: Molecular mechanisms of epithelial-mesenchymal transition. Nat Rev Mol Cell Biol 2014, 15(3):178–196. 10.1038/nrm3758

30. Voulgari A, Pintzas A: Epithelial–mesenchymal transition in cancer metastasis: mechanisms, markers and strategies to overcome drug resistance in the clinic. Biochim Biophys Acta 2009, 1796(2):75–90. 10.1016/j.bbcan.2009.03.002

31. Thiery JP: Epithelial-mesenchymal transitions in tumour progression. Nat Rev Cancer 2002, 2(6):442–454. 10.1038/nrc822

32. Wei SC, Fattet L, Tsai JH, Guo Y, Pai VH, Majeski HE, Chen AC, Sah RL, Taylor SS, Engler AJ et al: Matrix stiffness drives epithelial-mesenchymal transition and tumour metastasis through a TWIST1-G3BP2 mechanotransduction pathway. Nat Cell Biol 2015, 17(5):678–688. 10.1038/ncb3157

33. Hagihara Y, Okuzaki Y, Matsubayashi K, Kaneoka H, Suzuki T, Iijima S, Nishijima KI: Primordial germ cell-specific expression of eGFP in transgenic chickens. Genesis 2020, 58(8):e23388. 10.1002/dvg.23388

34. Yang J, Antin P, Berx G, Blanpain C, Brabletz T, Bronner M, Campbell K, Cano A, Casanova J, Christofori G: Guidelines and definitions for research on epithelial–mesenchymal transition. Nat Rev Mol Cell Biol 2020, 21(6):341–352. 10.1038/s41580-020-0237-9

35. Zeisberg M, Neilson EG: Biomarkers for epithelial-mesenchymal transitions. J Clin Invest 2009, 119(6):1429–1437. 10.1172/JCI36183

36. Radisky ES, Radisky DC: Matrix Metalloproteinase-Induced Epithelial-Mesenchymal Transition in Breast Cancer. J Mammary Gland Biol Neoplasia 2010, 15(2):201–212. 10.1007/s10911-010-9177-x

37. Potts JD, Runyan RB: Epithelial-mesenchymal cell transformation in the embryonic heart can be mediated, in part, by transforming growth factor β. Dev Biol 1989, 134(2):392–401. 10.1016/0012-1606(89)90111-5

38. Nawshad A, LaGamba D, Hay ED: Transforming growth factor beta (TGFbeta) signalling in palatal growth, apoptosis and epithelial mesenchymal transformation (EMT). Arch Oral Biol 2004, 49(9):675–689. 10.1016/j.archoralbio.2004.05.007

39. Gordon KJ, Kirkbride KC, How T, Blobe GC: Bone morphogenetic proteins induce pancreatic cancer cell invasiveness through a Smad1-dependent mechanism that involves matrix metalloproteinase-2. Carcinogenesis 2009, 30(2):238–248. 10.1093/carcin/bgn274

40. Xu Q, Wang L, Li H, Han Q, Li J, Qu X, Huang S, Zhao RC: Mesenchymal stem cells play a potential role in regulating the establishment and maintenance of epithelial-mesenchymal transition in MCF7 human breast cancer cells by paracrine and induced autocrine TGF-β. Int J Oncol 2012, 41(3):959–968. 10.3892/ijo.2012.1541

41. Domogatskaya A, Rodin S, Boutaud A, Tryggvason K: Laminin-511 but not-332,-111, or-411 enables mouse embryonic stem cell self-renewal in vitro. Stem Cells 2008, 26(11), 2800–2809. 10.1634/stemcells.2007-0389

42. Rodin S, Domogatskaya A, Ström S, Hansson EM, Chien KR, Inzunza J, Hovatta O, Tryggvason K: Long-term self-renewal of human pluripotent stem cells on human recombinant laminin-511. Nat Biotechnol 2010, 28(6), 611. 10.1038/nbt.1620

43. Anstrom KK, Tucker RP: Tenascin-C lines the migratory pathways of avian primordial germ cells and hematopoietic progenitor cells. Dev Dyn 1996, 206(4), 437–446. 10.1002/(sici)1097-0177(199608)206:4<437::Aid-aja9>3.0.Co;2-j

44. Biegler MT, Belay K, Wang W, Szialta C, Collier P, Luo J-D, Haase B, Gedman GL, Sidhu AV, Harter E et al: Pronounced early differentiation underlies zebra finch gonadal germ cell development. Dev Biol 2025, 517, 73–90. 10.1016/j.ydbio.2024.08.006

45. Nakano M, Arisawa K, Yokoyama S, Nishimoto M, Yamashita Y, Sakashita M, Ezaki R, Matsuda H, Furusawa S, Horiuchi H: Characteristics of Novel Chicken Embryonic Stem Cells Established Using Chicken Leukemia Inhibitory Factor. J Poult Sci 2011, 48(1):64–72. 10.2141/jpsa.010102

46. Livak KJ, Schmittgen TD: Analysis of relative gene expression data using real-time quantitative PCR and the 2(-Delta Delta C(T)) Method. Methods 2001, 25(4):402–408. 10.1006/meth.2001.1262

47. Bolger AM, Lohse M, Usadel B: Trimmomatic: a flexible trimmer for Illumina sequence data. Bioinformatics 2014, 30(15):2114–2120. 10.1093/bioinformatics/btu170

48. Martin M: Cutadapt removes adapter sequences from high-throughput sequencing reads. EMBnet journal 2011, 17(1):10–12. 10.14806/ej.17.1.200

49. Dobin A, Davis CA, Schlesinger F, Drenkow J, Zaleski C, Jha S, Batut P, Chaisson M, Gingeras TR: STAR: ultrafast universal RNA-seq aligner. Bioinformatics 2013, 29(1):15–21. 10.1093/bioinformatics/bts635

50. Li B, Dewey CN: RSEM: accurate transcript quantification from RNA-Seq data with or without a reference genome. BMC bioinformatics 2011, 12:1–16. 10.1186/1471-2105-12-323

51. Love MI, Huber W, Anders S: Moderated estimation of fold change and dispersion for RNA-seq data with DESeq2. Genome Biol 2014, 15(12):550. 10.1186/s13059-014-0550-8

52. Sun J, Nishiyama T, Shimizu K, Kadota K: TCC: an R package for comparing tag count data with robust normalization strategies. BMC Bioinformatics 2013, 14(1):219. 10.1186/1471-2105-14-219

53. Raudvere U, Kolberg L, Kuzmin I, Arak T, Adler P, Peterson H, Vilo J: g:Profiler: a web server for functional enrichment analysis and conversions of gene lists (2019 update). Nucleic Acids Res 2019, 47(W1):W191–w198. 10.1093/nar/gkz369

54. Zhou Y, Zhou B, Pache L, Chang M, Khodabakhshi AH, Tanaseichuk O, Benner C, Chanda SK: Metascape provides a biologist-oriented resource for the analysis of systems-level datasets. Nat Commun 2019, 10(1):1523. 10.1038/s41467-019-09234-6

55. Kolde R, Kolde MR: Package ‘pheatmap’. R package 2015, 1(7):790.

56. Kanda Y: Investigation of the freely available easy-to-use software ‘EZR’for medical statistics. Bone Marrow Transplant 2013, 48(3):452–458. 10.1038/bmt.2012.244

