## Supplementary_Figures for "A cell-matrix interaction regulates the undifferentiated state and self-renewal capacity of avian primordial germ cells"

### Slide 1
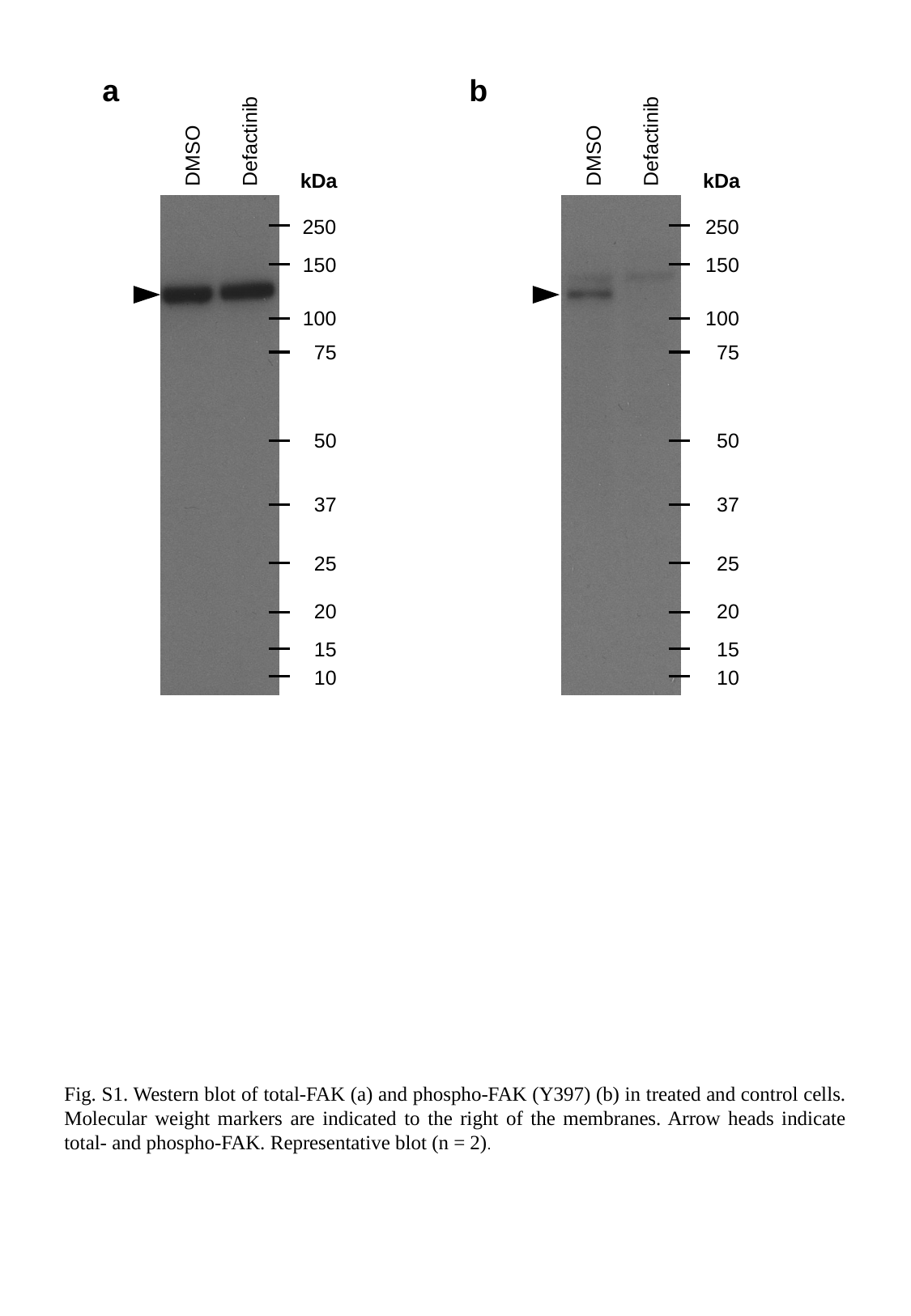

a
b
Defactinib
Defactinib
DMSO
DMSO
kDa
kDa
250
250
150
150
100
100
75
75
50
50
37
37
25
25
20
20
15
15
10
10
Fig. S1. Western blot of total-FAK (a) and phospho-FAK (Y397) (b) in treated and control cells. Molecular weight markers are indicated to the right of the membranes. Arrow heads indicate total- and phospho-FAK. Representative blot (n = 2).

### Slide 2
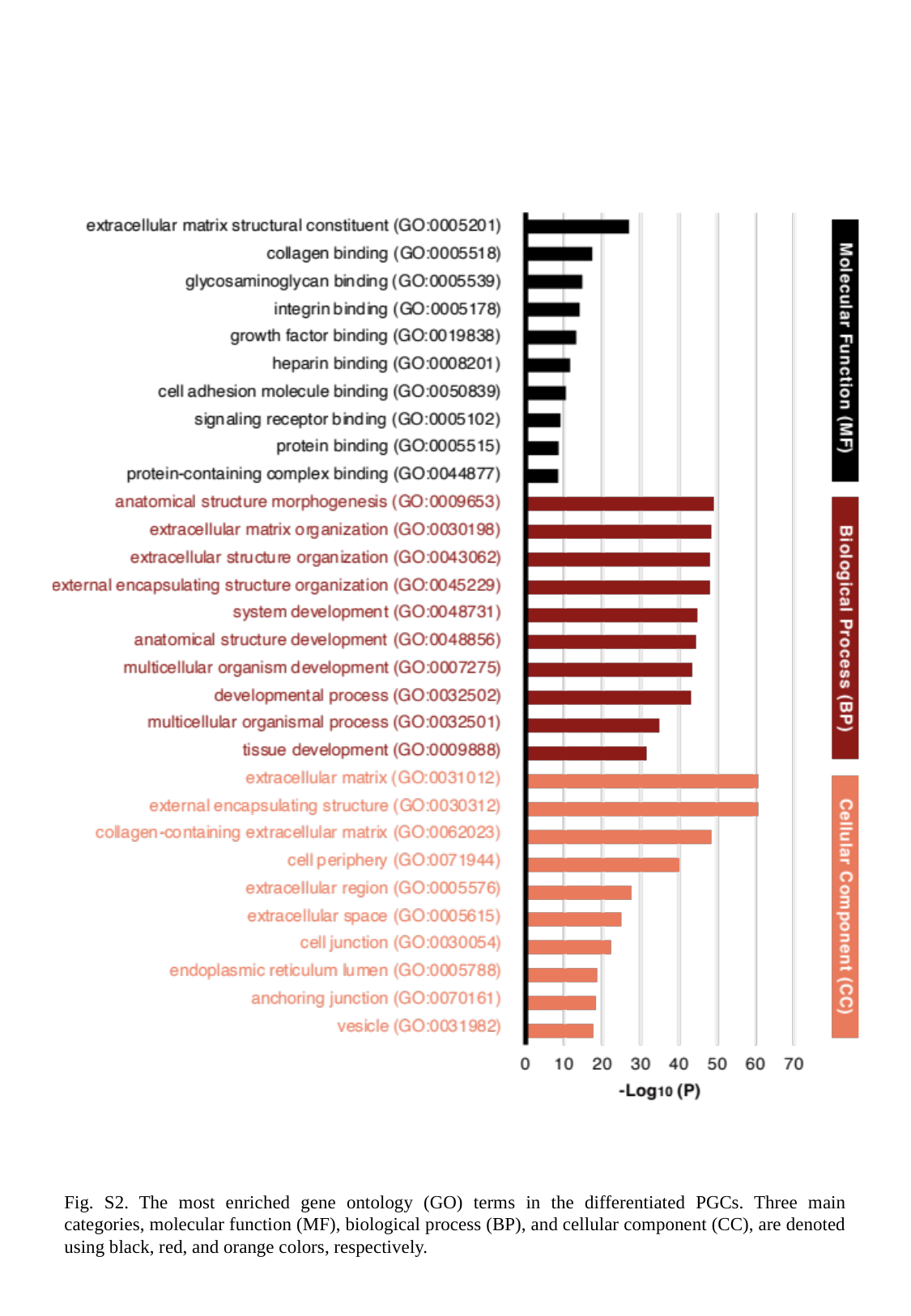

Fig. S2. The most enriched gene ontology (GO) terms in the differentiated PGCs. Three main categories, molecular function (MF), biological process (BP), and cellular component (CC), are denoted using black, red, and orange colors, respectively.

### Slide 3
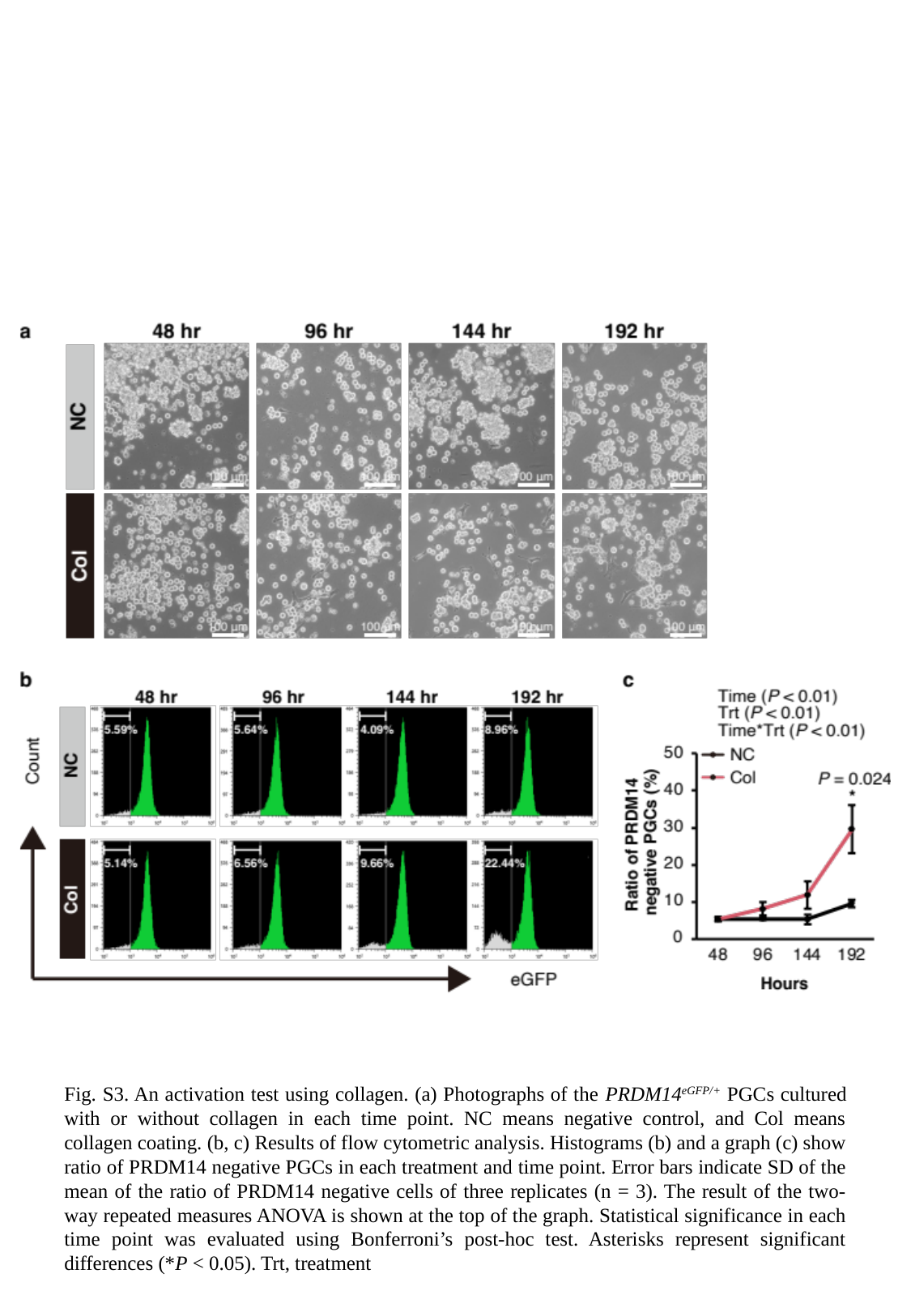

Fig. S3. An activation test using collagen. (a) Photographs of the PRDM14^eGFP/+^ PGCs cultured with or without collagen in each time point. NC means negative control, and Col means collagen coating. (b, c) Results of flow cytometric analysis. Histograms (b) and a graph (c) show ratio of PRDM14 negative PGCs in each treatment and time point. Error bars indicate SD of the mean of the ratio of PRDM14 negative cells of three replicates (n = 3). The result of the two-way repeated measures ANOVA is shown at the top of the graph. Statistical significance in each time point was evaluated using Bonferroni’s post-hoc test. Asterisks represent significant differences (*P < 0.05). Trt, treatment
